# Immunogenicity and pre-clinical efficacy of an OMV-based SARS-CoV-2 vaccine

**DOI:** 10.1101/2021.07.12.452027

**Authors:** Alberto Grandi, Michele Tomasi, Cinzia Bertelli, Teresa Vanzo, Assunta Gagliardi, Elena Caproni, Silvia Tamburini, Laura Fantappiè, Gabriele Di Lascio, Zeno Bisoffi, Chiara Piubelli, Maria Teresa Valenti, Luca Dalle Carbonare, Donato Zipeto, Micol Ravà, Valeria Fumagalli, Pietro Di Lucia, Davide Marotta, Eleonora Sala, Matteo Iannacone, Peter Cherepanov, Martino Bolognesi, Massimo Pizzato, Guido Grandi

**Author notes:** Corresponding authors: Guido Grandi, University of Trento, CIBIO Department, Via Sommarive 9, 28123, Trento Italy, Massimo Pizzato, University of Trento, CIBIO Department, Via Sommarive 9, 28123, Trento Italy.

## Abstract

The vaccination campaign against SARS-CoV-2 relies on the world-wide availability of effective vaccines, with a potential need of 20 billion vaccine doses to fully vaccinate the world population. To reach this goal, the manufacturing and logistic processes should be affordable to all countries, irrespectively of economical and climatic conditions.

Outer membrane vesicles (OMVs) are bacterial-derived vesicles that can be engineered to incorporate heterologous antigens. Given the inherent adjuvanticity, such modified OMVs can be used as vaccine to induce potent immune responses against the associated protein. Here we show that OMVs engineered to incorporate peptides derived from the receptor binding motif (RBM) of the spike protein from SARS-CoV-2 elicit an effective immune response in immunized mice, resulting in the production of neutralizing antibodies. The immunity induced by the vaccine is sufficient to protect K18-hACE2 transgenic mice from intranasal challenge with SARS-CoV-2, preventing both virus replication in the lungs and the pathology associated with virus infection. Furthermore, we show that OMVs can be effectively decorated with RBM peptides derived from a different genetic variant of SARS-CoV-2, inducing a similarly potent neutralization activity in vaccinated mice. Altogether, given the convenience associated with ease of engineering, production and distribution, our results demonstrate that OMV-based SARS-CoV-2 vaccines can be a crucial addition to the vaccines currently available.

## Introduction

The outbreak of SARS-CoV-2 has generated a pandemic event which caused almost 200 million documented infections worldwide and 4 million deaths to the current date^1^. Despite early attempts to contain the infection within the area where the first human cases were reported, the etiological agent of COVID-19 has spread worldwide and became the eighth human coronavirus. Unlike SARS-CoV and MERS, the current prevalence and the high rate of asymptomatic or pauci-symptomatic infections will favour SARS-CoV-2 circulation in adults and children in the future and will establish the infection as endemic, similarly to other human respiratory coronaviruses which cause flu-like pathologies (OC43, HKU1, 229E and NL63). Developing a long-term effective vaccination strategy is crucial to build the best possible immunity against SARS-CoV-2 in the human population, following the continuous evolution of the virus driven by the adaptation into the host and the selective pressure posed by the immune system. Reminiscent of the yearly influenza prophylaxis, the need of periodic updating and re-administration of the SARS-CoV-2 vaccine, depending on the extent of viral genetic drift, becomes a likely prospect for the future.

Currently, more than 280 candidate vaccines are under development at different stages, and, among these, 102 are in clinical phase ^2^. While vaccines based on protein subunits are reported to represent the largest fraction of those investigated, vaccines based on mRNA and viral vectors were the first to obtain emergency use authorization by the major agencies and were massively distributed and administered world-wide. As a result, the world witnessed an unprecedented technical and economical challenge, given the need of manufacturing, distributing, storing, and administering billions of doses of vaccines in every country, irrespectively of climatic, social and economic conditions. To overcome such challenges and provide a sustainable long-term prophylaxis, the vaccine production should rely on a process easily scalable and low-cost to make the doses affordable to the less favoured regions of the world. Importantly, to facilitate distribution and storage in all countries, the vaccine should not depend on the cold chain, which is impractical and logistically and economically prohibitive particularly in several parts of the world.

Among the several technologies available for vaccine development, outer membrane vesicles (OMVs) have emerged in recent years as an attractive tool capable of coupling excellent built-in adjuvanticity provided by the bacterial pathogen-associated-molecular patterns (PAMPs) associated with the vesicles, and an easily scalable production and purification process^3 4 5^. While OMVs purified from different pathogens are known to induce potent protective immunity against the same pathogens from which they are derived from (accordingly, anti-*Neisseria meningitidis* OMV-based vaccine are currently available for human use ^6 7 8^), we have recently developed a strategy which allows the engineering of *E. coli* OMVs selectively loaded with heterologous antigens ^9 10^. This strategy has been successfully demonstrated to induce potent immunity against foreign bacterial components and against human tumour-specific antigens. ^11,12^ Given that OMVs are readily phagocytosed, antigens carried by the vesicles are efficiently presented by professional antigen presenting cells, leading to efficient induction of a sustained Th1 response as well as an optimal humoral response. Since clinical evidence demonstrates that an accelerated induction of a Th1 cell response is associated with less severe cases of COVID-19^13,14^ and that convalescent individuals develop strong memory CD4+ and CD8+ T cells ^15^, the ability of OMVs to trigger Th1 represents a desired feature. Crucially, in addition to the simple and cost-effective setup required to produce and purify OMVs, the antigen-decorated vesicles are extremely stable for long-term storage at room temperature, making it a convenient vaccine to distribute all over the world^10^.

All SARS-CoV-2 vaccine candidates under development are designed to induce antibodies specific for the spike (S) protein. Neutralizing titres found in vaccinated individuals, as well as in convalescent patients, correlate strongly with antibody binding to the spike receptor binding domain (RBD), which interacts with the angiotensin-converting enzyme 2 (ACE2) on the host cell membrane ^16^. Accordingly, the most potent monoclonal antibodies isolated from convalescent patients recognize epitopes located in the RBD interface with ACE2^17^. The ability to specifically concentrate the immune response against these epitopes would therefore exclusively elicit neutralizing antibodies, while minimizing the risk of generating non-neutralizing or poorly neutralizing immunoglobulins binding to irrelevant spike regions. The OMVs offer the unique opportunity to display short and defined epitopes to B- and T-cells in a highly immunogenic context provided by the bacterial vesicle. Taking advantage of such potential and the availability of the crystal structure of the RBD in complex with ACE2, we engineered the OMVs with peptides derived from the RBM alpha helices which form the interface with ACE2. Here we show that RBM-derived peptides can be correctly associated with OMVs, inducing neutralizing antibodies titres sufficient to fully protect K18-hACE2 transgenic mice from challenge with SARS-CoV-2. Given the efficacy of the vaccine in the animal model, the ease of its engineering, the cost-effective production process, and the stability at room temperature, we propose the OMV-based vaccine as a promising candidate to continue the vaccination campaign against SARS-CoV-2.

## Results

### Design and construction of the OMV-based vaccine

As shown by the 3D structure of the SARS-CoV-2 RBD in complex with ACE2, the concave receptor binding motif (RBM) of the spike protein is organized in two discrete ordered chains incorporating the β5 and β6 strands which cross each other and include most of the residues contacting ACE2 (Figure 1A). Patient-derived monoclonal antibodies (mAbs) binding epitopes in these chains of the RBD potently neutralize virus cell entry *in vitro* and are currently in clinical use or in advanced clinical development ^18^. In the attempt of producing a vaccine eliciting neutralizing immunity against SARS-CoV-2, we generated OMVs decorated with the polypeptides derived from the RBM, fused with FhuD2, a *Staphylococcus aureus* lipoprotein shown to efficiently deliver heterologous protein domains to the *E. coli* outer membrane and to the vesicular compartment^10^.

**Figure 1.**
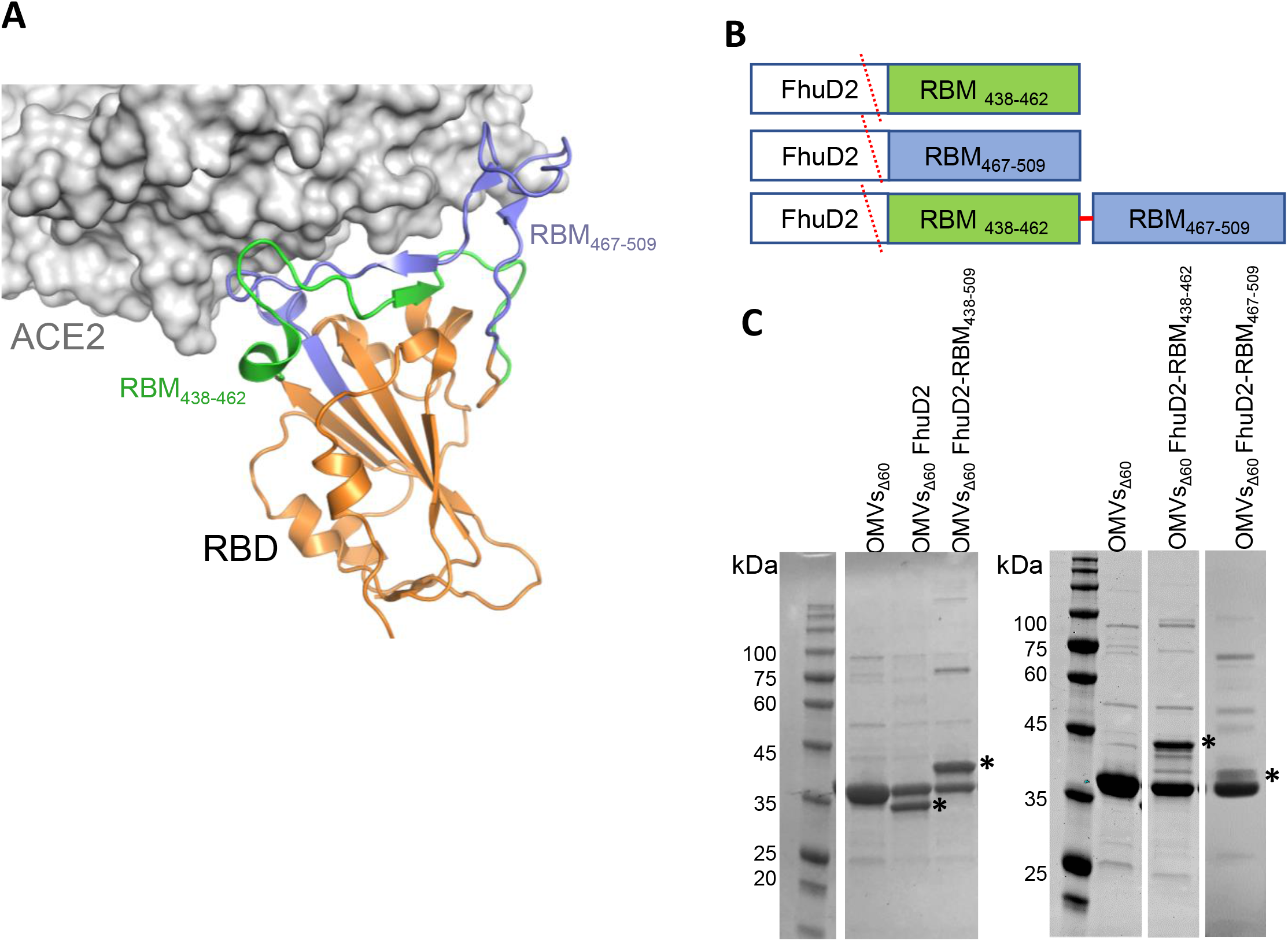
Construction and production of OMVs carrying SARS-CoV-2 RBM antigens. (A) Topology of the interaction between SARS-CoV-2 RBD and ACE2 with the indication of the two RBM polypeptides tested in this study. (B) Schematic representation of the pET21-based constructs expressed in BL21(DE3)*Δ60* to decorate OMVs. *Staphylococcus aureus* Ferric hydroxamate receptor 2 (FhuD2) was fused to one copy of RBM_438-462_, RBM_467-509_ and RBM_438-509_ SARS-CoV-2 epitopes. (C) SDS-PAGE of purified OMVs derived from BL21(DE3)*Δ60* cultures expressing pET21-FhuD2-RBM_438-462_, pET21-FhuD2-RBM_467-509_ and pET21-FhuD2-RBM_438-509_ plasmids or the control pET21-FhuD2 or empty pET21 plasmids. Stars indicate the predicted migration of the fusion proteins.

To this end, the nucleotide sequence coding for β5 (RBM_438-462_) and β6 (RBM_467-509_) strands or a combination of the two (RBM_438-509_) were fused to the 3’ end of the sequence encoding FhuD2 in plasmid pET-FhuD2^12^, thus generating the fusion proteins FhuD2-RBMs (Figure 1B). The resulting plasmids were used to transform *E. coliΔ60*, a hyper-vesiculating *E. coli* BL21(DE3) derivative recently created in our laboratories^10^.

From the culture supernatant of one recombinant clone each (*E. coliΔ60(*pET-FhuD2-RBMs)) OMVs were concentrated and purified. In order to verify the correct association of the fusion proteins with the bacterial vesicles, FhuD2-RBMs-OMVs were subjected to SDS-PAGE analyses, which revealed the presence of protein species compatible with the predicted molecular weights of the FhuD2-RBM fusion proteins (Figure 1C). Based on the intensity of the bands visualized by Coomassie staining, the incorporation of the fusion protein into OMVs appears to be similar to the incorporation level of FhuD2 alone, indicating that the addition of the SARS-CoV-2 RBM polypeptides is well tolerated and compatible with efficient transport into the bacterial vesicles.

### Immunogenicity of FhuD2-RBMs-OMVs

Having proven efficient association with OMVs, we next tested the capacity of FhuD2-RBMs-OMVs to induce the production of antibodies capable of recognizing the RBD in the context of the SARS-CoV-2 spike protein. To this aim, five CD1 mice were immunized with each construct three times at two-week intervals with 10 μg of each engineered OMV preparation combined with aluminum hydroxide (Alum) (2 mg/ml, Figure 2A) administered intraperitoneally. Mouse sera were collected seven days after the third dose from each immunization group, pooled, and the level of IgGs binding the SARS-CoV-2 RBD domain were detected by ELISA (Figure 2B) with plates coated with the spike RBD domain expressed and purified from HEK293T cells. As shown in Figure 2, all sera of animals immunized with the different FhuD2-RBMs-OMVs contained high amounts of specific IgGs recognizing the RBM (Figure 2B), with optical deviation (OD) 1 obtained with dilutions between 1:7000 and 1:30000 of the sera.

**Figure 2.**
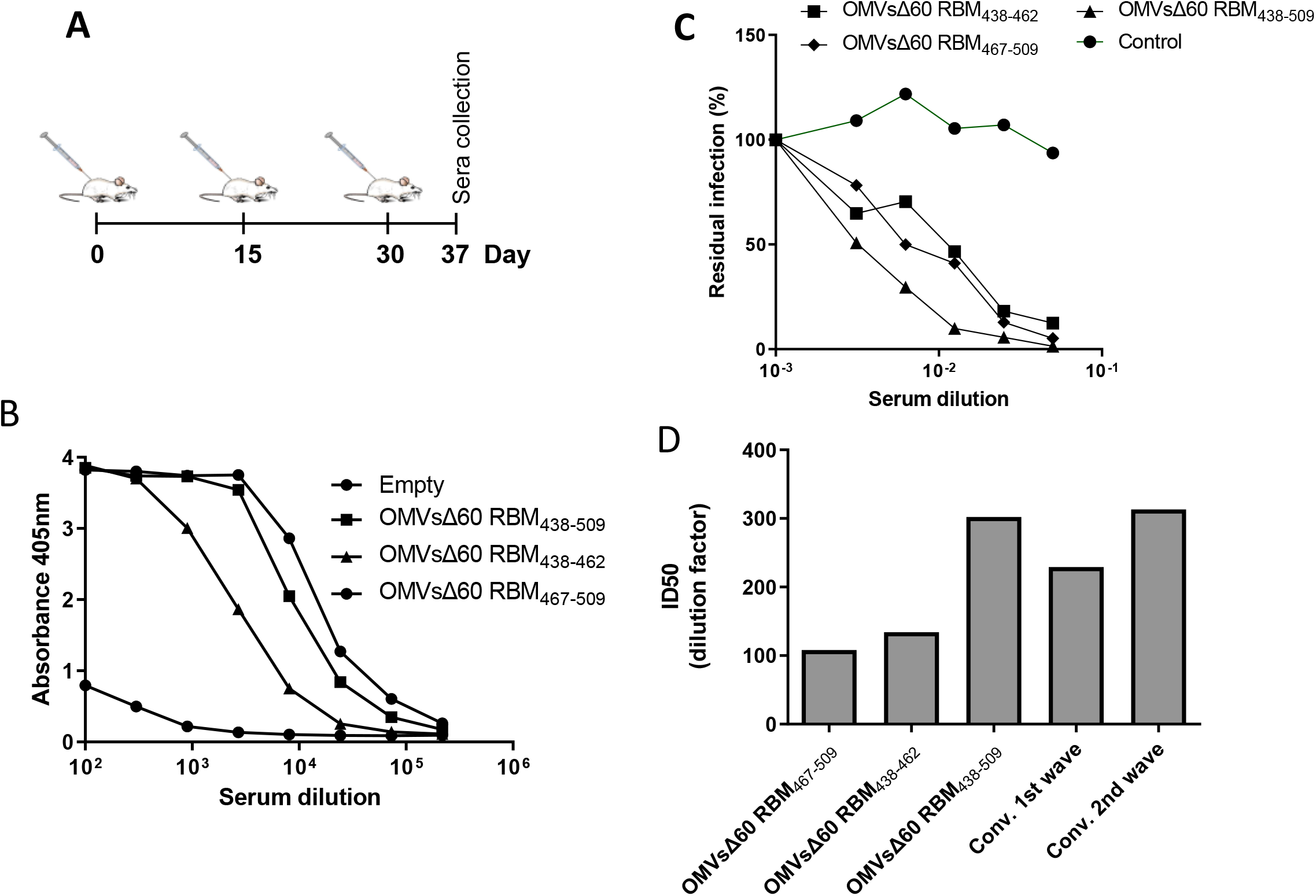
Immunogenicity and of OMVs decorated with SARS-CoV-2 RBM antigens in immunized mice. (A) Diagram of the experimental setup of the immunization experiments. CD1 mice received three intraperitoneal immunizations on day 0, 14 and 28 with 10 μg of OMVs together with 2mg/ml Aluminium hydroxide. Sera were collected 10 days after the last immunization. As control, mice were immunized with non-engineered OMVs. (B) Antibody titers in sera pooled from each group of 5 mice, collected 10 days after the third immunization, measured by ELISA, using plates coated with the SARS-CoV-2 RBD. (C) Neutralization activity in pooled sera, measured with lentiviral vectors pseudotyped with SARS-CoV-2 spike from the Whuan-1 isolate, plated on Huh-7 cells. Residual infectivity after treatment with serially diluted sera, expressed as percentage of the untreated virus control is shown. (D) Comparison of neutralization activity measures in pooled sera from groups of immunized animal (5 animal per group) and in convalescent individuals who contracted the infection with SARS-CoV-2 in spring 2020 (first wave, 25 individuals) or in autumn 2020 (second wave, 25 individuals). The histogram shows TCID50 values derived from (C) and from analyses performed with the same neutralization assay using sera from the two convalescent groups (average values are shown).

Having demonstrated the presence of high titers of anti-spike antibodies in mice immunized with OMVs incorporating all three fusion proteins, we next asked whether such antibodies could also neutralize virus infection *in vitro*. To this aim, neutralization was assessed with a pseudovirus assay using a GFP- encoding SIV-based lentiviral vector pseudotyped with the SARS-CoV-2 Spike protein derived from the Wuhan-1 isolate. Sera derived from each group of mice immunized with the engineered OMVs were pooled, serially diluted and combined with the pseudotyped vectors before inoculation on Huh-7-ACE2 target cells.

The data shown in Figure 2C demonstrate that the sera collected from mice immunized with all three different OMVs incorporating the RBM fragments neutralize SARS-CoV-2 spike pseudotyped vectors. While the OMVs incorporating the single strands (RBM_438-462_ and RBM_438-569_) neutralized with similar potency with an ID50 of 1:108 and 1:134 respectively, OMVs associated with the combination of the two β strands (RBM_438-569_) induced a more powerful neutralization (ID50 of 1:302), indicating that both chains contribute and synergize to the elicitation of neutralizing antibodies.

In order to compare the neutralization induced in mice vaccinated with RBM OMVs with the titers elicited by the natural infection in the human population, we used the same pseudovirus assay to analyze the neutralizing titers in 50 convalescent individuals who contracted the infection during either the first or the second COVID-19 pandemic waves. We found that the neutralizing titers measured in individuals 8-10 months after contracting COVID-19 (first wave) displayed in average ID50 of 229 while the titers observed in individuals that contracted the infection more recently (2-4 months, during the second wave) was in average 313. The neutralization activity elicited by the RBM_438-569_ OMVs in mice is therefore of similar magnitude compared with the neutralization induced by the natural infection observed in convalescent patients.

### Protective activity of FhuD2-RBM_438-509_-OMVs in hACE2 transgenic mice

Having established that the RBMs OMVs are effective at inducing neutralizing immunity in mice, and that vesicles which incorporate the combination of two RBM strands elicit higher immunity compared with those displaying either the β5 or β6 strands alone, we selected the former to explore the degree of protection induced *in vivo* against SARS-CoV-2 challenge. To this end, K18-hACE2 transgenic mice19 received two intraperitoneal immunizations (at day −28 and at day −14) of 10 μg of FhuD2-RBM_438-509_ OMVs vaccine (*n* = 4) or PBS (*n* = 4) (Fig. 3A). Fourteen days after the boost immunization, all mice were infected intranasally with 1 × 10^5^ TCID_50_ of SARS-CoV-2 (hCoV-19/Italy/LOM-UniSR-1/2020; GISAID Accession ID: EPI_ISL_413489) (Fig. 3A). As expected^20^, beginning 4 days post infection (p.i.) PBS-treated K18-hACE2 transgenic mice infected with SARS-CoV-2 developed a severe disease, as assessed by monitoring respiration, coat condition, posture, social behavior, and palpebral aperture^21^ (Fig. 3B). By contrast, K18-hACE2 transgenic mice immunized with the FhuD2-RBM_438-509_ OMVs vaccine developed a much milder disease (Fig. 3B). Viral RNA was measured (Fig. 3C) and infectious virus was titrated (Fig. 3D) from lungs of infected mice 5 days after challenge. Out of four vaccinated mice, only in one mouse viral RNA and replicating virus could be detected, indicating the ability of the vaccine to prevent virus replication in the respiratory tract. Mirroring the relative abundance of viral RNA, the N protein was also readily detected by both immunohistochemistry (Fig. 3E) and immunofluorescence microscopy (Fig. 3F) in the lungs of PBS-treated mice while it was mostly absent in mice immunized with the FhuD2-RBM_438-509_ OMVs. The effective protection by the vaccine should also reflect the absence or attenuation of immune activation in response to the viral challenge. To assess the degree of inflammation triggered by the viral infection, we measured the amount of inflammatory monocytes (CD11b+Ly6C high and CD64+, figures 3G and H respectively) recruited in the lungs after infection. In vaccinated mice a significantly lower number of inflammatory monocytes were recovered from the lungs of vaccinated mice compared to unvaccinated animals after viral challenge, correlating with the lower viral titers (Fig. 3 C, D, E, F).

**Figure 3.**
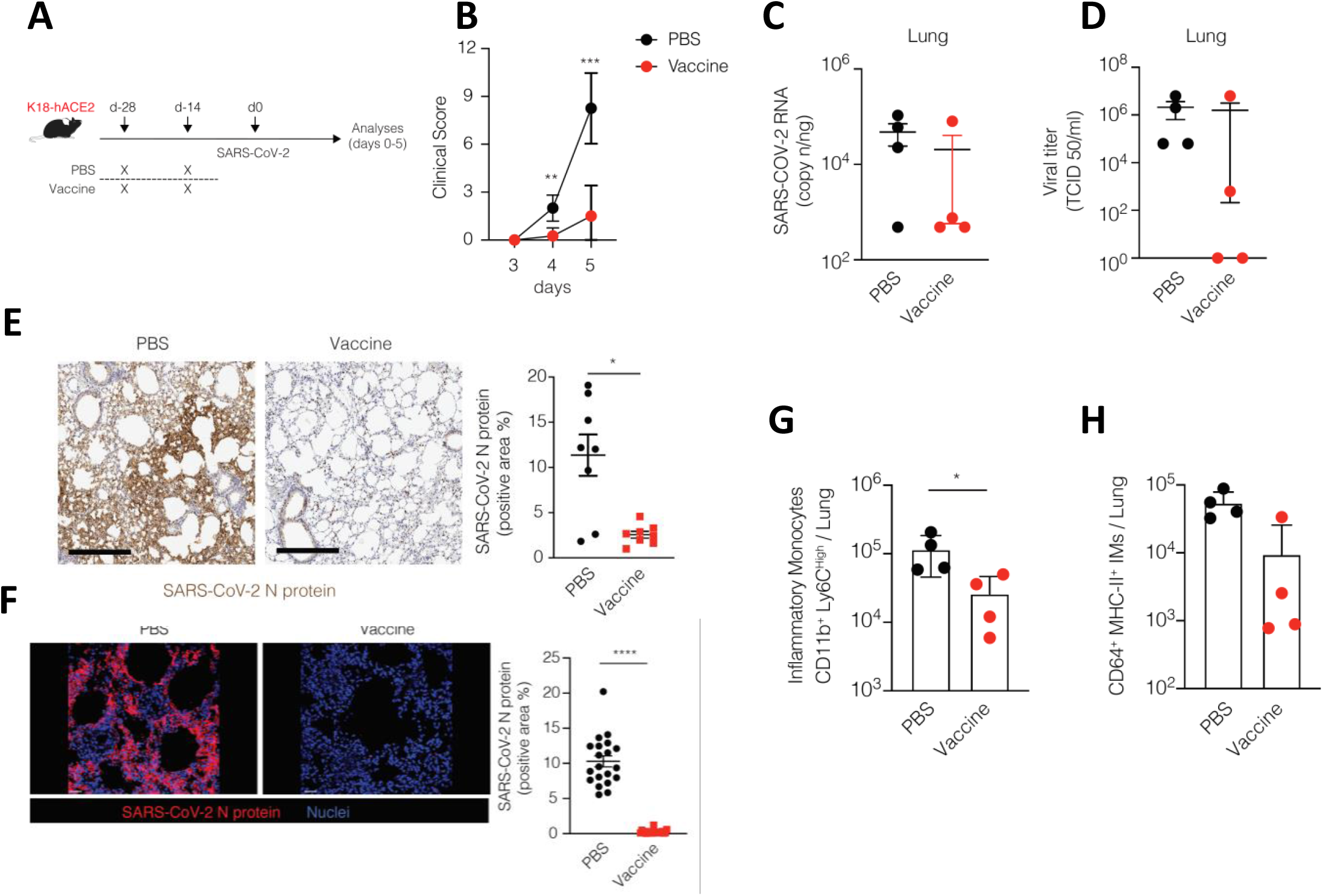
*In vivo* protection efficacy against SARS-CoV-2 challenge in hACE2 transgenic mice. (A) Schematic representation of the experimental setup. K18-hACE2 mice (C57BL/6 background) received two intraperitoneal immunizations (at day - 28 and - 14) of 10 μg of OMV vaccine (n = 4) or PBS (n = 4) prior to intranasal infection with 1×10^5^ TCID50 of SARS-CoV-2. Lung were collected and analyzed five days after SARS-CoV-2 infection. (B) Mice were observed daily for clinical symptoms to assess severity of disease based on respiration, coat condition, posture, social behavior, and palpebral aperture. (C) SARS-CoV-2 RNA in the lung was quantified by quantitative PCR with reverse transcription (RT–qPCR) 5 days after infection. (D) Viral titers in the lung 5 days after infection were determined by median tissue culture infectious dose (TCID50). (E) Representative immunohistochemical micrographs of lung sections 5 days post SARS-CoV-2 infection. N-SARS-CoV-2 expression is shown in brown. Scale bars, 300 μm. Right panel, quantification of N-SARS-CoV-2 signal, each dot represents a mouse. (F) Representative confocal immunofluorescence micrographs of lung sections from PBS-treated mice (left) or immunized mice (right) 5 days post SARS-CoV-2 infection. N-SARS-CoV-2 positive cells are depicted in red and nuclei in blue. Scale bar represents 30 μm. Right panel, quantification of N-SARS-CoV-2 signal, each dot represents a different stack. (G) Absolute numbers of CD11b+ Ly6Chigh inflammatory monocytes in the lung of the indicated mice 5 days after SARS-Cov-2 infection. (H) Absolute numbers of CD64+ MHC-II+ inflammatory monocytes in the lung of the indicated mice. * p value < 0.05, ** p value < 0.01, *** p value < 0.001, **** p value < 0.0001.

### Elicitation of neutralizing antibodies with OMVs decorated with the RBM_438-509_ epitope from the P.1 isolate

In order to test the adaptability of the OMV-based vaccine to RBM from different SARS-CoV-2 variants, we engineered the vesicles using the combination of β5 and β6 strands derived from the P.1 variant (B.1.1.28.1) which carries two changes (E484K, N501Y) in the RBM compared to the Wuhan-1 isolate (Fig. 4A). The peptide, fused to FhuD2, could be expressed in *E. coli* and was associated efficiently to OMVs, as observed by SDS-PAGE (Fig. 4B). Mice were immunized with the engineered OMVs following the time schedule described in fig 2A and sera collected 7 days after the last inoculation. High amount of IgG interacting with the RBD were elicited by the vaccine, as assessed by ELISA on RBD (Figure 4C). Of note, while ELISA plates were coated with RBD derived from the Wuhan-1 isolate, the amount of IgGs in the animal sera capable of binding the RBD was similar to that induced by OMV decorated with the RBM derived from the Wuhan-1 isolate (Figure 4C), despite the presence of two polymorphic amino acids. Moreover, sera collected from mice immunized with OMVs decorated with the RBM_438-509_ from the P.1 variant neutralized SARS-CoV-2 Wuhan-1 spike pseudotyped vectors with similar potency respect to sera from mice immunized with OMVs engineered with the RBM_438-509_ epitope from the Wuhan-1 isolate (Figure 4D). In conclusion, our results prove that the OMVs can be successfully engineered to induce neutralizing immunity with RBM form different SARS-CoV-2 variants.

**Figure 4.**
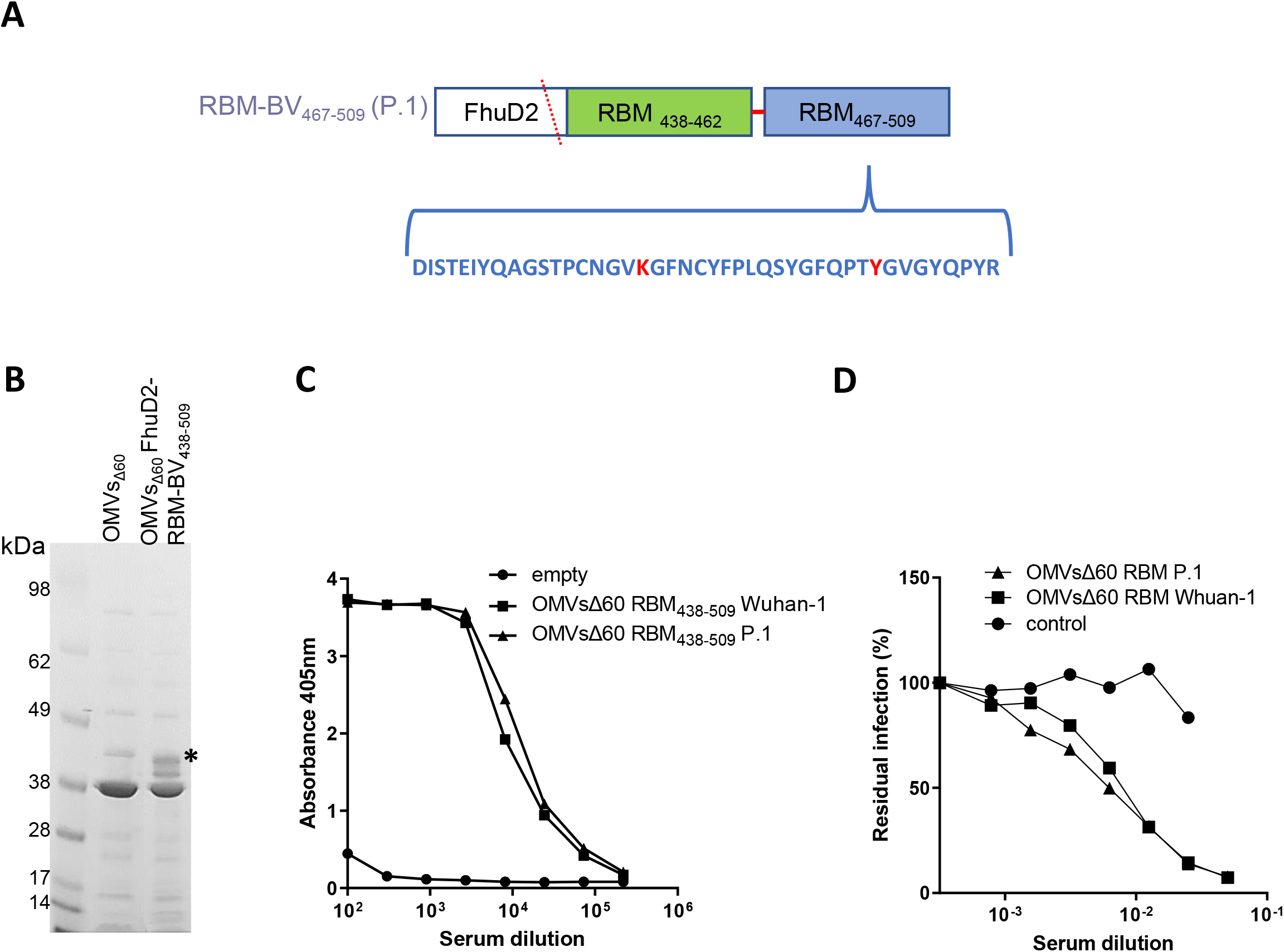
Production and immunogenicity of OMVs decorated with RBM antigens derived from the P.1 isolate of SARS-CoV-2. (A) Schematic representation of the the pET21-based construct expressed in BL21(DE3)*Δ60* to decorate OMVs. FhuD2 was fused to the RBM_438-509_ SARS-CoV-2 epitope derived from the P.1 isolate. The sequence shows in red the two RBM amino acids which differ between the Wuhan-1 and the P.1 isolates (E484K and N501K). CD1 mice received three intraperitoneal immunizations on day 0, 14 and 28 with 10 μg of OMVs together with 2mg/ml Aluminium hydroxide. Sera were collected 10 days after the last immunization. As control, mice were immunized with non-engineered OMVs. (B) SDS-PAGE of purified OMVs derived from BL21(DE3)*Δ60* cultures expressing pET21-FhuD2-P.1-RBM_438-509_ plasmids or the control empty pET21 plasmid. Stars indicate the predicted migration of the fusion protein. (C) Antibody titers in sera pooled from each group of 5 mice, collected 10 days after the third immunization, measured by ELISA, using plates coated with the SARS-CoV-2 RBD. (C) Neutralization activity in pooled sera from each group of 5 mice, measured with lentiviral vectors pseudotyped with SARS-CoV-2 spike from the Whuan-1 isolate, plated on Huh-7 cells. Residual infectivity after treatment with serially diluted sera, expressed as percentage of the untreated virus inoculum control is shown.

## Discussion

Despite the plethora of anti-SARS-CoV-2 vaccines under development and the availability of a few extremely effective vaccines, the effort to offer additional prophylactic solutions to the COVID-19 pandemic is still worthy of consideration for a number of reasons. First, to guarantee the completion of the global vaccination campaign in an acceptable timeframe, more than 20 billion vaccine doses should be available as soon as possible. Such massive amount of vaccine doses can only be provided with the contribution of several vaccines developed at different production sites distributed worldwide. At present, we are still far from this ambitious objective. As a matter of fact, many countries, particularly low-income countries, have just started their vaccination campaigns and it is difficult to predict if, when and how such campaigns can be completed. Second, the difficulty in providing sufficient vaccines doses to all population is also associated to the costs of vaccine production and distribution. Available vaccines require sophisticated technologies and production plants, which are difficult to reproduce in different sites. Moreover, the vaccines require storage conditions with difficult and expensive logistical solutions. Third, the duration of immunological protection guaranteed by both natural infection and vaccination remains unclear. A possible picture which is emerging is the need of an annual administration of booster doses of vaccine in order to provide protection and prevent viral spreading. Should this be the case, the issue of vaccine production shortage would be further amplified. Finally, the need of booster doses might be further exacerbated by the appearance of novel viral variants which might require modified versions of the vaccines to be fully neutralized.

With this complex scenario, we believe that an OMV-based vaccine could offer interesting advantages and solutions. First of all, OMVs appear amenable to display crucial peptides of the Spike glycoprotein. Of note, while properly folded eukaryotic glycoproteins can normally only be expressed in a eukaryotic expression system, requiring a labor-intensive protein purification process, the crucial portion of the spike RBM, which contacts directly ACE2, can be efficiently expressed and incorporated into OMVs while maintaining a natural conformation. An effective immunity is in fact elicited in vaccinated mice, with efficient production of neutralizing antibodies, at levels sufficient to protect the animals from infection. Accordingly, the reduced viral replication in the lungs of vaccinated mice was associated with a favorable clinical score and a severely reduced recruitment of inflammatory monocytes.

Second, OMVs are extremely easy to produce. The separation of the biomass from the culture supernatant and an ultrafiltration step to concentrate OMVs and eliminate contaminants released into the supernatant is essentially all is needed for vaccine production^22^. The process is amenable to large scale production and can be easily transferred to different sites to expand vaccine production. Moreover, the production yields make the vaccine costs particularly affordable. Under laboratory conditions, we reproducibly obtain more than 5.000 OMV-based vaccine doses/liter of culture. Practically speaking, it means that using a 1.000-liter fermentation unit associated to a tangential flow ultrafiltration devise is sufficient to provide 5 million doses of vaccine/week at costs that are expected to be well below 1 USD/dose.

Third, although not yet experimentally demonstrated, theoretically speaking our OMV vaccine should be particularly indicated as a booster vaccine both in the case the boost is needed to reinvigorate the immune response against the original vaccine strain and in the case the immune response has to be potentiated against a new viral variant. Accordingly, OMVs can be easily and quickly modified to incorporate updated antigens.

In the first case, as the data here presented demonstrate, our vaccine has the unique property to elicit antibodies which specifically bind the RBM region interacting with the ACE2 receptor. Therefore, when the immune response goes below a protective threshold, it could be convenient to boost those memory B and T cells which are particularly relevant for protection. This would avoid the “dilution” of the response toward irrelevant epitopes. Moreover, the potent adjuvanticity of the OMVs should further enhance the optimal expansion of functional B and T cells.

As far as the use of a booster dose to cope with the spreading of new variants is concerned, theoretically speaking our OMV-based vaccine should mitigate the risk of the so call “original antigenic sin (OAS)”. OAS refers to the fact that once the immune system is exposed to a certain pathogen, a second infection by a slightly different version of the same pathogen preferentially triggers the immunological memory generated by the first infection^23^. As a consequence, the regions of the pathogen which differ in the new variant are poorly recognized by the immune system, with the risk that the immune responses are not sufficient to neutralize the new variant. Thanks to the fact that our OMV vaccine specifically targets the key domains necessary and sufficient to neutralize the virus, the OAS effect should be avoided.

In conclusion given the convenience associated with the engineering, production and storage, the efficacy as a vaccine against SARS-CoV-2 makes *E. coliΔ60* OMVs a valuable tool for the future management of COVID19.

## Materials and Methods

### Engineering BL21(DE3) E. coli strains with SARS-CoV-2 neutralizing epitopes

The pET21-FhuD2-RBM_438-462_, pET21-FhuD2-RBM_467-509_ and pET21-FhuD2-RBM_438-509_ plasmids carrying the *Staphylococcus aureus* Ferric hydroxamate receptor 2 (FhuD2) fused to one copy of RBM_438-462_, RBM_467-509_ and RBM_438-509_ SARS-CoV-2 epitopes respectively were assembled using the PIPE method as described in (ref. H. E. Klock, S. A. Lesley, The Polymerase Incomplete Primer Extension (PIPE) method applied to high-throughput cloning and site-directed mutagenesis. *Methods Mol. Biol.* **498**, 91–103 (2009)). Briefly, pET21-FhuD2 plasmid was linearized by PCR, using FhuD2-v-R and pET-V-F primers (Table 1). In parallel, the synthetic DNA encoding the RBD of SARS-CoV-2 was synthesized by GeneArt (Thermo Fisher Scientific, Waltham, MA, USA) and used as template for the amplification of the three RBM epitopes (Table 2). More in detail, RBM_438-462_ and RBM_467-509_ were amplified by PCR with the forward 2-F and 1-F and the reverse 2-R and 1-R primers, respectively (Table 1). The PCR products and the linearized plasmid were mixed together and used to transform *E. coli* HK100 strain.

**Table 1.**
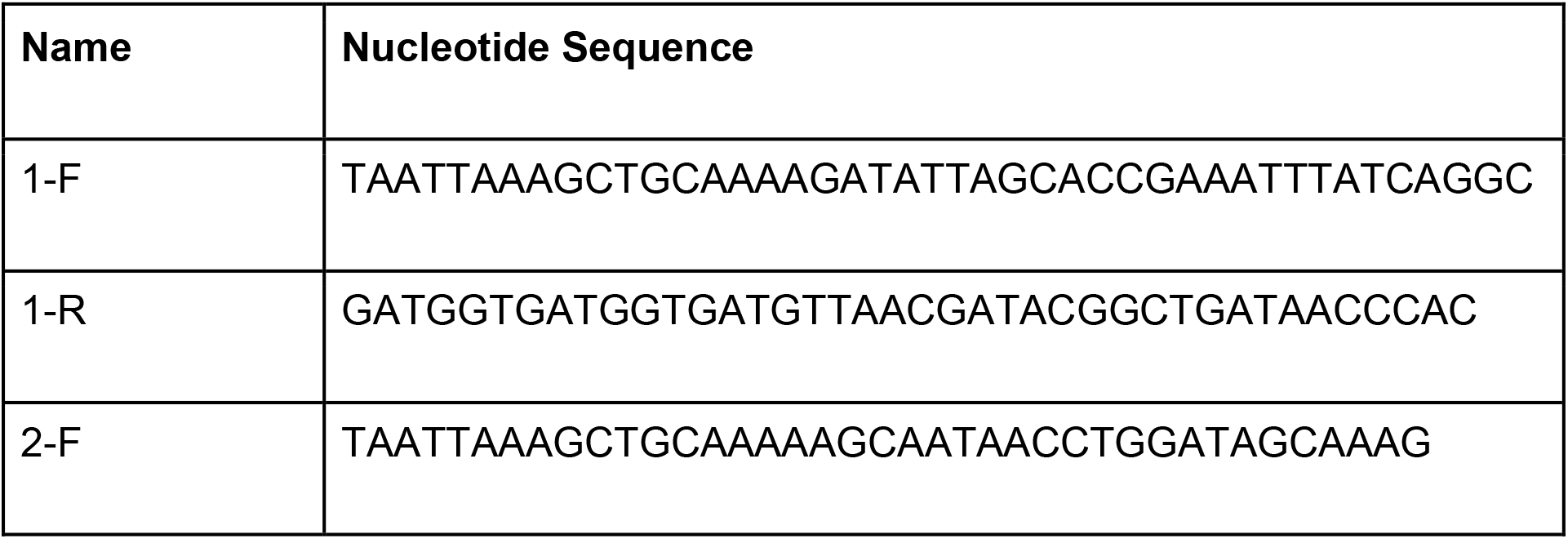
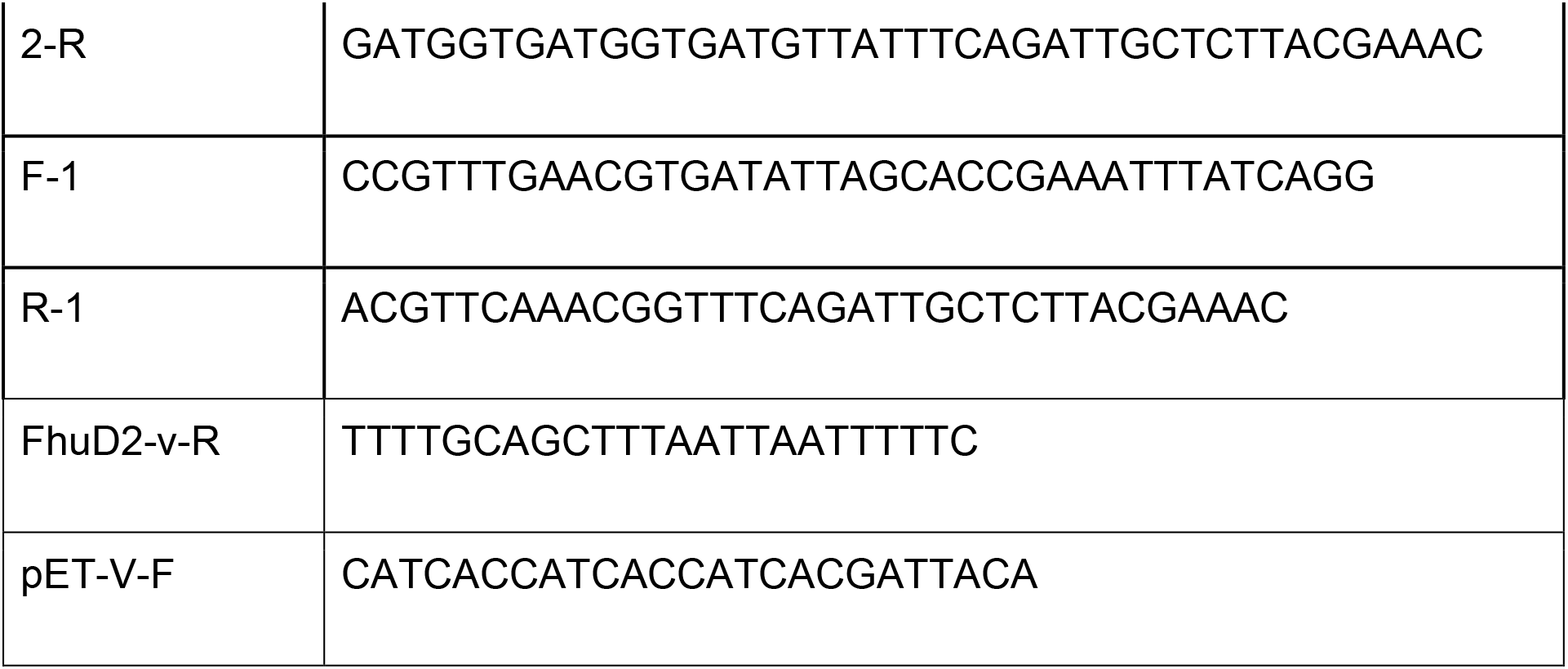
(oligo)

**Table 2.**
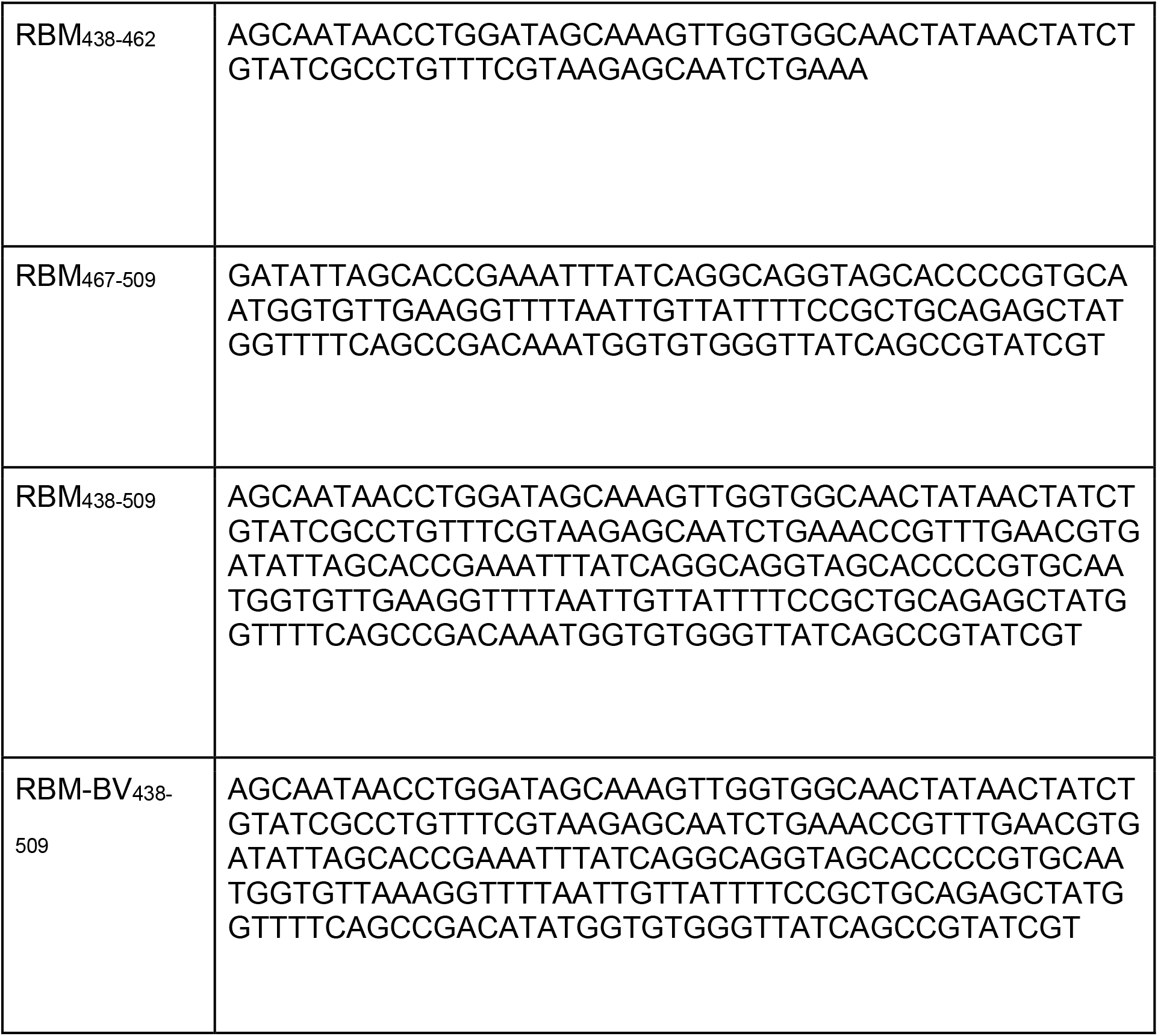
(sequences)

RBM_438-509_, the combination of the two epitopes, was assembled with two steps of PCR in succession. In the first step two different fragments carrying an overlapping sequence were amplified with 2-F/R-1 and F-1/1-R primers (Table 1). In the second step the two fragments were eluted from Agarose gel, mixed together and used as template for a final PCR reaction with the primers 2-F and 1-R. This final product and the linearized plasmid were mixed together and used to transform the *E. coli* HK100 strain.

The RBM-BV438-509 gene, encoding for the RBM_438-509_ epitope from the P.1 isolate, a variant of the RBD carrying the E484K and N501Y mutations, was synthesized by GeneArt (Thermo Fisher Scientific, Waltham, MA, USA) and used as template for the amplification of the RBM-BV_438-509_ using the 2-F and 2-R primers. The PCR amplification product and the pET21-FhuD2 linearized plasmid were mixed together and used to transform *E. coli* HK100 strain.

To confirm the correctness of the gene fusions, plasmids were sequenced (Eurofins, Ebersberg, Germany, EU) and *E. coli* BL21(DE3)Δ60 strain was transformed with pET21-FhuD2-RBM_438-462_, pET21-FhuD2-RBM_467-509_, pET21-FhuD2-RBM_438-509_ and pET21-FhuD2-RBM-BV_438-509_ plasmids and the derived recombinant strains were used for the production of engineered FhuD2-RBM_438-462_, FhuD2-RBM_467-509_, FhuD2-RBM_438-509_ and FhuD2-RBM-BV_438-509_ OMVs, respectively.

### OMV purification

OMVs from BL21(DE3)*Δ60*(pET-FhuD2-RBM_438-509_), BL21(DE3) *Δ60*(pET-FhuD2-RBM_438-462_), BL21(DE3)*Δ60*(pET-FhuD2-RBM_467-509_) and BL21(DE3)*Δ60*(pET-FhuD2-RBM-BV_438-509_), were purified in an EZ control bioreactor (Applikon Biotechnology, Schiedam, Netherlands) as previously described (Zanella et al., 2021). Cultures were started at an OD_600_ of 0.1 and grown in LB medium at 30°C, pH 6.8 (±0.2), dO_2_ > 30%, 280–500 rpm until OD_600_=0.5, then the temperature was lowered to 25°C. The expression of the recombinant protein was induced when the culture reached an OD_600_ of 1 with 0.1 mM IPTG and a feed made of 50 mg/l ampicillin, 15 g/l glycerol, 0.25 g/l MgSO_4_ was added. The fermentation was carried out until the end of the exponential phase at 25°C. OMVs were then purified and quantified as previously described (Zanella et al., 2021). Culture supernatants were separated from biomass by centrifugation at 4000g for 20 minutes. After filtration through a 0.22-μm pore size filter (Millipore, Burlington, Massachusetts, USA), OMVs were isolated, concentrated and diafiltrated from the supernatants using Tangential Flow Filtration (TFF) with a Cytiva Äkta Flux system. OMVs were quantified using DC protein assay (Bio-Rad, Hercules, California. USA).

### SDS-PAGE analysis

20 μg (protein content) were resuspended in sodium dodecyl sulfate-polyacrylamide gel electrophoresis (SDS-PAGE) Laemmli buffer and heated at 98°C for 10’. Proteins were separated using Any kD™ Criterion™ TGX Stain-Free™ Protein Gel (BioRad) in TrisGlicyne buffer (BioRad) and revealed by Coomassie Blue staining.

### Enzyme-Linked Immunosorbent Assay

Ninety six-well Maxisorp plates (Nunc, Thermo Fisher Scientific) were coated overnight with 200 ng/well of purified recombinant RBD in PBS at 4°C. The following day, the plate was blocked with 100 ml/well of PBS+1% BSA for 30 minutes at room temperature. Mice sera were threefold serially diluted in PBS+1% BSA starting from a 1:100 initial dilution. After 3 washes with PBS+0.05% Tween 20 (PBST), 100 μl of each serum dilution were added in each well and the plate was incubated at room temperature for 1 hour. Wells were washed three times with PBST and then incubated 45 min at room temperature with goat anti mouse alkaline phosphatase-conjugate antibodies at a final dilution of 1:2000 (SigmaAldrich). After 3 washes with PBST, 100 ml/well of 3 mg/ml paranitrophenyl-phosphate disodium hexahydrate (Sigma Aldrich) in 1 M diethanolamine buffer pH 9.8 and plates were incubated at room temperature in the dark for 45min. Finally, absorbance was read at 405 nm using Varioskan™ LUX multimode microplate reader.

### Animal studies

Mice were monitored twice per day to evaluate early signs of pain and distress, such as respiration rate, posture, and loss of weight (more than 20%) according to humane endpoints. Animals showing such conditions were anesthetized and subsequently sacrificed in accordance with experimental protocols, which were reviewed and approved by the Animal Ethical Committee of The University of Trento and the Italian Ministry of Health.

Five-week old CD1 female mice were immunized intraperitoneally (i.p.) on day 0, 14 and 28 with 10 μg of OMVs together with 2mg/ml Aluminium hydroxide. Sera were collected 10 days after the last immunization.

For virus challenge studies, B6.Cg-Tg(K18-ACE2)2Prlmn/J mice1 were purchased from The Jackson Laboratory. Mice were housed under specific pathogen-free conditions and heterozygous mice were used at 8-10 weeks of age. All experimental animal procedures were approved by the Institutional Animal Committee of the San Raffaele Scientific Institute and all infectious work was performed in designed BSL-3 workspaces.

The hCoV-19/Italy/LOM-UniSR-1/2020 (EPI_ISL_413489) isolate of SARS-CoV-2 was obtained from the Unit of Microbiology and Virology of San Raffaele Scientific Institute and grown in Vero E6 cells.

K18-hACE2 transgenic mice were immunized intraperitoneally with 10μl vaccine or PBS twice, 14 days apart. Virus infection was performed via intranasal administration of 1 × 105 TCID50 SARS-CoV-2 per mouse.

### Measurement of viral titers from infected animals

Tissues homogenates were prepared by homogenizing perfused lung using gentleMACS Octo Dissociator (Miltenyibiotec, #130-096-427) in M tubes (#130-093-335) containing 1 ml of DMEM. Samples were homogenized for three times with program m_Lung_01_02 (34 seconds, 164 rpm). The homogenates were centrifuged at 3’500 rpm for 5 minutes at 4°C. The supernatant was collected and stored at −80°C for viral isolation and viral load detection.

Viral titers were calculated by 50% tissue culture infectious dose (TCID_50_). Briefly, Vero E6 cells were seeded at a density of 1.5 × 10^4^ cells per well in flat-bottom 96-well tissue-culture plates. The following day, 2-fold dilutions of the homogenized tissue were applied to confluent cells and incubated 1 h at 37°C. Then, cells were washed with phosphate-buffered saline (PBS) and incubated for 72 h at 37°C in DMEM 2% FBS. Cells were fixed with 4% paraformaldehyde for 20 min and stained with 0.05% (wt/vol) crystal violet in 20% methanol.

### RNA extraction and qPCR

Tissues homogenates were prepared by homogenizing perfused lung using gentleMACS dissociator (Miltenyibiotec, #130-096-427) with program RNA_02 in M tubes (#130-096-335) in 1 ml of Trizol (Invitrogen, #15596018). The homogenates were centrifuged at 2000 g for 1 min at 4°C and the supernatant was collected. RNA extraction was performed by combining phenol/guanidine-based lyisis with silica membrane-based purification. Briefly, 100 μl of Chloroform were added in 500 ml of homogenized sample and total RNA was extracted using ReliaPrep™ RNA Tissue Miniprep column (Promega, Cat #Z6111). Total RNA was isolated according to the manufacturer’s instructions. qPCR was performed using TaqMan Fast virus 1 Step PCR Master Mix (Lifetechnologies #4444434), standard curve was drawn with 2019_nCOV_N Positive control (IDT#10006625), primer used are: 2019-nCoV_N1-Forward Primer (5’-GAC CCC AAA ATC AGC GAA AT-3’), 2019-nCoV_N1-Reverse Primer (5’-TCT GGT TAC TGC CAG TTG AAT CTG-3’) 2019-nCoV_N1-Probe (5’-FAM-ACC CCG CAT TAC GTT TGG TGG ACC-BHQ1-3’) (Centers for Disease Control and Prevention (CDC) Atlanta, GA 30333). All experiments were performed in duplicate.

### Cell Isolation and Flow Cytometry

Mice were euthanized by cervical dislocation. Lungs were perfused through the right ventricle with PBS at the time of autopsy. Lung tissue was digested in RPMI 1640 containing 3.2 mg/ml Collagenase IV (Sigma, #C5138) and 25 U/ml DNAse I (Sigma, #D4263) for 30 minutes at 37°C. Homogenized lungs were passed through 70 μm nylon mesh to obtain a single cell suspension. Cells were resuspended in 36% percoll solution (Sigma #P4937) and centrifuged for 20 minutes at 2000 rpm (light acceleration and low brake). The remaining red blood cells were removed with ACK lysis.

Cell viability was assessed by staining with Viobility™ 405/520 fixable dye (Miltenyi, Cat #130-109-814). Antibodies (Abs) used are indicated in the table below.

Flow cytometry analysis were performed on a BD FACSymphony A5 SORP and analyzed with FlowJo software (Treestar).

### Confocal immunofluorescence histology and histochemistry

Lungs of infected mice were collected and fixed in 4% paraformaldehyde (PFA). Samples were then dehydrated in 30% sucrose prior to embedding in OCT freezing media (Bio-Optica). 20 μm sections were cut on a CM1520 cryostat (Leica) and adhered to Superfrost Plus slides (Thermo Scientific). Sections were then permeabilized and blocked in PBS containing 0.3% Triton X-100 (Sigma-Aldrich) and 5% FBS followed by staining in PBS containing 0.3% Triton X-100 and 1% FBS. Slides were stained for SARS-CoV-2 nucleocapsid (GeneTex) for 1h RT. Then, slides were stained with Alexa Fluor 568 Goat Anti-Rabbit antibody for 2h RT. All slides were analyzed by confocal fluorescence microscopy (Leica TCS SP5 Laser Scanning Confocal). For N-SARS-CoV-2 immunohistochemistry, mice were perfused with PBS and lung were collected in Zn-formalin and transferred into 70% ethanol 24 h later. Tissue was then processed, embedded in paraffin and automatically stained for SARS-CoV-2 (2019-nCoV) Nucleocapsid Antibody (SINO BIO, 40143-R019) through LEICA BOND RX 1h RT and developed with Bond Polymer Refine Detection (Leica, DS9800). Brightfield images were acquired through an Aperio Scanscope System CS2 microscope and an ImageScope program (Leica Biosystem) following the manufacturer’s instructions. In both immunofluorescence and histochemistry, N-SARS-CoV-2 percentage of positive area was determined by QuPath (Quantitative Pathology & Bioimage Analysis) software.

### Clinical score

Mice were observed daily for clinical symptoms. Disease severity was scored as reported in Table 4.

**Table 3.** (antibodies).

| Name | Clone | Source and catalog number |
| --- | --- | --- |
| CD103 | 2E7 | Biolegend #121407 |
| MHCII I-a/I-b | M5/114.15.2 | Biolegend #107622 |
| Ly-6C | HK1.4 | Biolegend #128026 |
| CD11b | M1/70 | Thermo Fisher #48-0112-82 |
| SIGLEC F | E50-2440 | BD Biosciences #740388 |
| CD64 | X54-5/7.1 | BD OptiBuild #740622 |
| CD8 | 53-6.7 | Biolegend #100759 |
| CD11c | HL3 | BD Biosciences #563735 |
| CD4 | RM4-5 | BD Biosciences #740208 |
| CD44 | IM7 | BD Biosciences #741227 |
| CD69 | H1.2F3 | BD Biosciences #612793 |
| F4/80 | BM8 | Biolegend #123110 |
| Ly6g | 1A5 | BD Pharmingen #562700 |
| CD45 | 30-F11 | Biolegend #103113 |

**Table 4.**
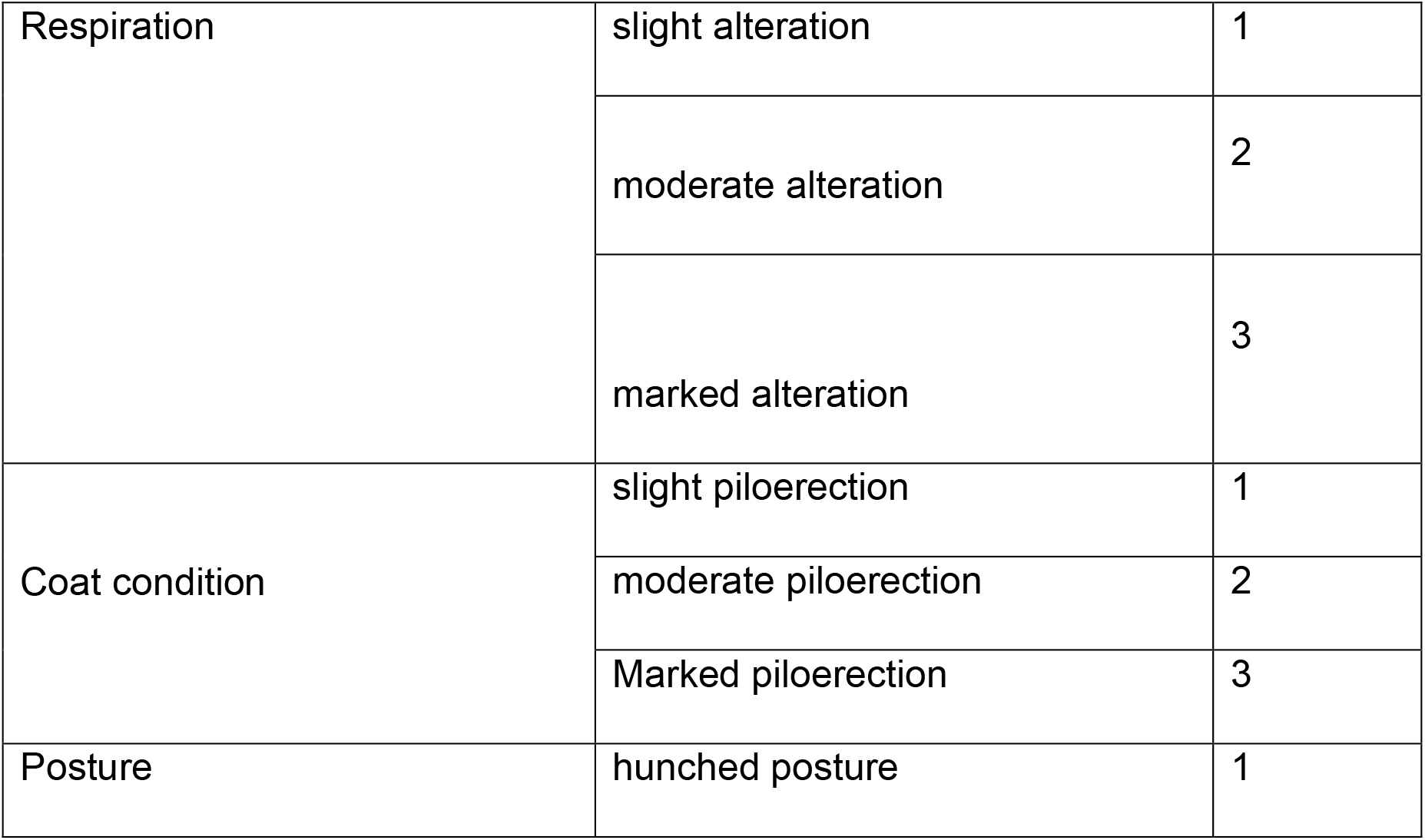
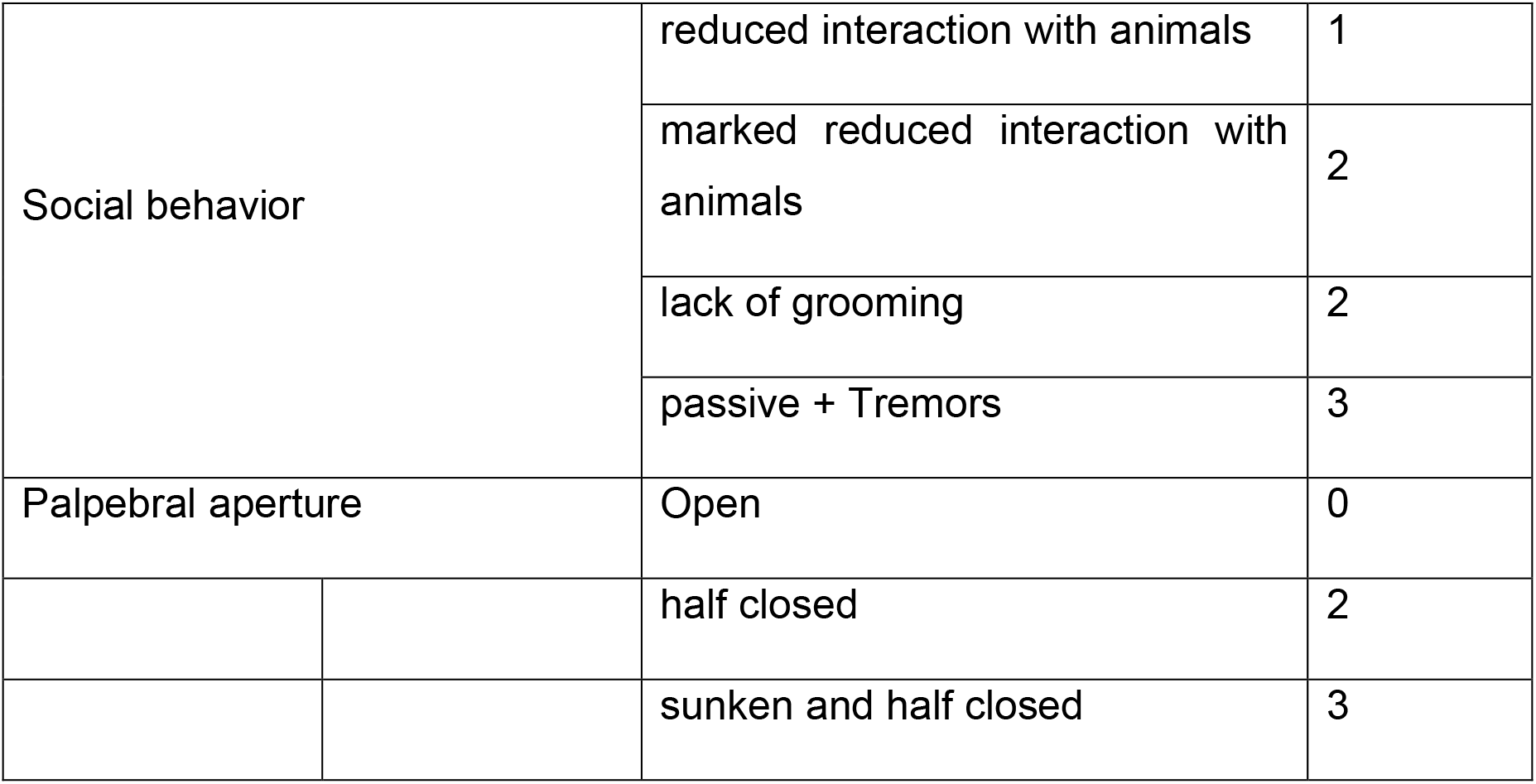
(clinical scores)

### Preparation of viral pseudotypes and neutralization assays

SIV-based lentiviral particles pseudotyped with SARS-Cov-2 spike were produced in 10 cm plates prepared the day before transfection with 3 million HEK293T cells in 10 ml complete DMEM, supplemented with 10% FBS. Cells were transfected using the Calcium Phosphate method with 15 μg of env-defective SIV-Mac239-GFP construct with GFP expressed in place of Nef (PMID 26181333) and 1.5 μg of PCDNA3.1 encoding the WT SARS-CoV-2 spike (reference sequence Wuhan-Hu-1, accession number YP_009724390) engineered to truncate the C-terminal 19 amino acids. Pseudotyped vector supernatants were harvested 48 h post-transfection and filtered through a 0.45-μm filter before. Neutralization titres were tested on Huh-7 cells overexpressing ACE2. Target cells were seeded onto 96-well tissue culture one day before neutralization. The vectors inoculum was adjusted to produce no more than 10% transduction of the monolayer to ensure a linear working range of the assay.

Sera dilutions were added to pseudotyped virus particles, incubated at room temperature for 30 minutes and added to cells. After 48 h, transduction was assessed by counting the GFP-expressing cells using the fluorescent plate reader EnSight (Perkin Elmer). Each serum dilution was assessed in triplicate. Neutralization was measured by calculating the residual transduction activity of the pseudovirus considering the untreated sample as 100%. Fitted sigmoidal curves and IC50 were obtained using Prism (GraphPad) with the least square variable slope method and using the dose-normalized response protocol.

### Human sera

The sera of 50 healthcare workers with pre-existing immunity for SARS-CoV-2 (as per former nasal swab positivity) were collected in January 2021. Samples had been collected and stored in the University of Verona biobank (Ethics Committee approval prot. N. 1538) and in Tropica Biobank of the IRCCS Sacro Cuore Don Calabria Hospital (Ethics Committee approval prot. N. 50950). All participants signed informed consent.

### Statistical analyses and software

Flow data were collected using FlowJo Version 10.5.3 (Treestar). Statistical analyses were performed with GraphPad Prism software version 8 (GraphPad). *n* represents individual mice analyzed per experiments. Error bars indicate the standard error of the mean (SEM). We used Mann-Whitney U-tests to compare two groups with non-normally distributed continuous variables. Significance is indicated as follows: *p<0.05; **p<0.01; ***p<0.001; ****p<0.0001. Comparisons are not statistically significant unless indicated.

## Acknowledgments

We thank E. Bono, L. Giustini and M. Mainetti for technical assistance. Flow cytometry was carried out at FRACTAL, a flow cytometry resource and advanced cytometry technical applications laboratory established by the San Raffaele Scientific Institute. Confocal immunofluorescence histology was carried out at Alembic, an advanced microscopy laboratory established by the San Raffaele Scientific Institute and the Vita-Salute San Raffaele University. We would like to acknowledge the PhD program in Basic and Applied Immunology and Oncology at Vita-Salute San Raffaele University, as D.M. and E.S. conducted this study as partial fulfillment of their PhD in Molecular Medicine within that program. M.I. is supported by the European Research Council (ERC) Consolidator Grant 725038, ERC Proof of Concept Grant 957502, Italian Association for Cancer Research (AIRC) Grants 19891 and 22737, Italian Ministry of Health Grants RF-2018-12365801 and COVID-2020-12371617, Lombardy Foundation for Biomedical Research (FRRB) Grant 2015-0010, the European Molecular Biology Organization Young Investigator Program, and Funded Research Agreements from Gilead Sciences, Takis Biotech and Asher Bio.

## Footnotes

Patent EP20175072.8 (BACTERIAL OUTER MEMBRANE VESICLES CARRYING CORONAVIRUS PROTEINS, METHOD OF PREPARATION, COMPOSITIONS AND USE THEREOF) submitted by BiOMViS on May 2020.

